# A microfabricated, 3D-sharpened silicon shuttle for insertion of flexible electrode arrays through dura mater into brain

**DOI:** 10.1101/614685

**Authors:** Hannah R. Joo, Jiang Lan Fan, Supin Chen, Jeanine A. Pebbles, Hexin Liang, Jason E. Chung, Allison M. Yorita, Angela C. Tooker, Vanessa M. Tolosa, Charlotte Geaghan-Breiner, Demetris K. Roumis, Daniel F. Liu, Razi Haque, Loren M. Frank

## Abstract

Electrode arrays for chronic implantation in the brain are a critical technology in both neuroscience and medicine. Recently, flexible, thin-film polymer electrode arrays have shown promise in facilitating stable, single-unit recordings spanning months in rats. While array flexibility enhances integration with neural tissue, it also requires removal of the dura mater, the tough membrane surrounding the brain, and temporary bracing to penetrate the brain parenchyma. Durotomy increases brain swelling, vascular damage, and surgical time. Insertion using a bracing shuttle results in additional vascular damage and brain compression, which increase with device diameter; while a higher-diameter shuttle will have a higher critical load and more likely penetrate dura, it will damage more brain parenchyma and vasculature. One way to penetrate the intact dura and limit tissue compression without increasing shuttle diameter is to reduce the force required for insertion by sharpening the shuttle tip. We describe a novel design and fabrication process to create silicon insertion shuttles that are sharp in three dimensions and can penetrate rat dura, for faster, easier, and less damaging implantation of polymer arrays. Sharpened profiles are obtained by reflowing patterned photoresist, then transferring its sloped profile to silicon with dry etches. We demonstrate that sharpened shuttles can reliably implant polymer probes through dura to yield high quality single unit and local field potential recordings for at least 95 days. On insertion directly through dura, tissue compression is minimal. This is the first demonstration of a rat dural-penetrating array for chronic recording. This device obviates the need for a durotomy, reducing surgical time and risk of damage to the blood-brain barrier. This is an improvement to state-of-the-art flexible polymer electrode arrays that facilitates their implantation, particularly in multi-site recording experiments. This sharpening process can also be integrated into silicon electrode array fabrication.

## Introduction

Electrode arrays for implantation in the brain are a technology critical to both fundamental neuroscience and clinical treatments for diseases including epilepsy (Toth et al., 2016), retinal degeneration (Luo and da Cruz, 2016), Parkinson’s, and depression (Lozano et al., 2019). Classically, electrode arrays have been made of silicon (D. Kipke, 2003, Rousche and Normann, 1998) or other hard metal (Nicolelis et al., 2003). While these devices can be effective in recording single units (Mols et al., 2017), the longevity of recordings is limited (Polikov et al., 2005, Jeong et al., 2015). More recent designs of silicon electrodes with smaller cross sections can record an estimated two cells per electrode, and can detect single units in mouse for at least 120 days (Jun et al., 2017), but the ability to record continuously from a given neuron over days has yet to be demonstrated.

Silicon electrodes are typically stiff, inspiring the development of flexible neural implants including probes (Chung et al., 2019, Jeong et al., 2015, Fu et al., 2016, Sohal et al., 2014, Xie et al., 2015) and mesh (Zhou et al., 2017) that are better matched mechanically to brain tissue and show reduced inflammation and immune reponses (Harris et al., 2011, Szarowski et al., 2003, Lee et al., 2017, Lacour et al., 2016, Jeong et al., 2015, Hong and Lieber, 2019), particularly if they are also small (Seymour and Kipke, 2007, Luan et al., 2017, Kozai et al., 2012). Chronically implanted in animal models, polymer devices can yield single-cell recordings spanning months (Chung et al., 2019, Jeong et al., 2015, Fu et al., 2016, Luan et al., 2017), with many of the same individual neurons recorded continuously for at least ten days (Chung et al., 2019).

The same flexibility that is desirable for a device once in the brain makes its implantation challenging. The brain and spinal cord are encased in three layers of dense irregular connective tissue, the dura, arachnoid, and pia mater (the meninges), that together protect the brain from mechanical and other insults (Weller et al., 2018). While some stiff devices can be implanted in monkey or rat brain after removal of the dura, a flexible device cannot typically penetrate even the remaining pia. This remains a major barrier to use for many flexible devices (Lecomte et al., 2018).

Solutions to the problem of inserting a flexible electrode array through rat pia for chronic recording include temporarily stiffening the array during insertion by associating it with biodissolvable coatings of silk (Wu et al., 2015), maltose (Xiang et al., 2014), or polyethylene glycol (PEG; Patel et al., 2015, Khilwani et al., 2016, Kozai et al., 2014, Lo et al., 2015, Kim et al., 2013), or filling a microfluidic channel within the device (Takeuchi et al., 2005). Other solutions include syringe injection (Liu et al., 2015, Schuhmann et al., 2018) or, to reduce the amount of fluid injected to the brain parenchyma, fluidic microdrive insertion (Vitale et al., 2018). Devices have also been frozen for insertion (Xie et al., 2015), or coated with adjustable shape memory polymer (SMP; Simon et al., 2017), which can penetrate dura when combined with an insertion guide (Shoffstall et al., 2018). One straightforward solution is to temporarily attach the flexible array to a hard shuttle, such as silicon (Kozai and Kipke, 2009, Felix et al., 2012, Chen et al., 2017, Patel et al., 2015), diamond (Na et al., 2018), or tungsten (Zhao et al., 2019) that is shape-matched to the device itself and can be retracted once the array has been inserted to its target depth in the brain. Temporary attachment can be achieved by threading electrodes through shuttles (Hanson et al., 2019) or, more commonly, by a biodissolvable adhesive. An advantage to this general method is that a very stiff material can be used to fabricate the insertion shuttle, since it will not be left in the brain. The shuttle diameter can thus be smaller, and the paired device very flexible, for reduced damage to the tissue on implantation and over the lifetime of a chronic implant (Seymour and Kipke, 2007). Previously, we have used such a method to insert 16 independent polymer devices through rat pia, each to a different brain area, for high-quality, chronic recording (Chung et al., 2019).

For a shuttle made of a given material, with length determined by the depth of the target brain area, a lower limit on the allowable device diameter is calculable by Euler’s critical load equation. A larger diameter device will have a higher critical load to more likely penetrate the meninges, but is not desirable because it will compress a greater area of brain tissue on insertion and disrupt more vasculature (Obaid et al., 2018). One way to enable insertion of a device without increasing its diameter is to reduce the effective force on the device (*i.e*., to lower the required insertion force) by fabricating a sharper tip (Sharp et al., 2009, Bjornsson et al., 2006, Edell et al., 1992, Jensen et al., 2006). If such a device could penetrate not only the brain but the encasing dura, then the tissue damage and longer surgical time that result from durotomy itself would also be eliminated. Furthermore, a dural-penetrating shuttle could also avoid errors in depth targeting that result from outward swelling of the brain following durotomy. We therefore aimed to develop a low-diameter sharpened shuttle that could penetrate dura with minimal brain compression.

Here we demonstrate that a silicon shuttle sharpened in three dimensions reliably penetrates dura. Using this shuttle, we report successful implantation and recording of local field potential (LFP) and single units from devices targeted to the orbitofrontal cortex (OFC), for at least 95 days. Although promising strategies for insertion of flexible polymer devices through rat dura have recently been demonstrated (Shoffstall et al., 2018, Zhao et al., 2019), such methods have not yet yielded chronic, high-quality recordings in freely behaving animals. This is the first *in vivo* demonstration of successful chronic neural recording using a method to implant through a membrane with Young’s modulus in the range 0.1-1 mPa (Maikos et al., 2008), corresponding to the rat dura or primate pia. 3D-sharpened shuttles obviate the need for a durotomy, increasing the efficiency and reliability of insertion for polymer electrode arrays to that of dural-penetrating arrays while maintaining the desirable properties of polymer for high quality single-unit neural recordings.

## Materials and methods

We fabricated a sharpened shuttle for insertion of flexible, thin-film polymer probes and tested transdural insertion *in vivo* in the rat. *In vivo* tests and chronic recording were performed in 10-16 month-old Long Evans rats. To evaluate this method, we measured insertion force and brain compression, which have been shown to predict neural tissue damage (Dixon et al., 1991). First, we compared the insertion force through intact dura for shuttles that were either sharpened in three dimensions (sharpened) or two dimensions (planar) with otherwise identical dimensions. We first tested insertion through dura over the hippocampus, then tested insertion through the thicker dura over the OFC. Second, we calculated the brain compression in each case. Finally, we implanted polymer devices using sharpened shuttles through intact dura over the nucleus accumbens (NAc) and OFC, and recorded LFP and single units for 95 days.

### 1.1 Shuttle microfabrication

Planar and sharpened shuttles both had a 30 degree tip angle (Fig. 1), a diameter of 80 µm, and a thickness of 30 µm (Fig. 2). The difference in their design was the gradual increase in thickness (the “sharpened” profile) of the 3d-sharpened shuttle (Fig. 3). Both planar and sharpened insertion shuttles were fabricated on 4” silicon-on-insulator (SOI) wafers with 30 µm device layers (Fig. 4). First, 10 µm deep “wicking” channels (Fig. 1, Fig. 2) were fabricated using standard photolithography and the Bosch Deep Reactive Ion Etch (DRIE) process in an STS ICP DRIE tool (SPTS Technologies). Wicking channels were designed to hold polyethylene glycol (PEG, molecular weight 10,000 mn), which acted as a biodissolvable adhesive to mount polymer probes on insertion shuttles, as described in Felix *et al.*, 2012. Next, 300 nm of aluminum was sputtered (Semicore Equipment Inc.), patterned photolithographically, and wet etched to create a hard mask with the insertion shuttle geometry. Sharpened shuttle fabrication involved a third photolithography step to specify the sharpened tip profile overlaid on the aluminum hard mask, followed by a 2-minute reflow bake of the photoresist (AZ4620) at 180 degrees Celsius (Fig. 4 a, b). This resulted in a sloped profile in the patterned photoresist that was transferred to silicon using alternating timed silicon etch and photoresist etch steps (Fig 4 c, d, e) in the same STS etcher. The silicon DRIE process used C_4_F_8_ for temporary sidewall protection and SF_6_ for etching. The photoresist etch process used only O_2_ for etching. Different silicon profiles could be achieved from the same reflowed photoresist profile by tailoring the alternating etch times. Planar shuttles underwent DRIE without the third photolithography step, resulting in a tip profile of uniform thickness. Lastly, the aluminum hard mask was removed (Piranha etch) and individual shuttles were released from the SOI wafer using hydrofluoric acid.

**Figure 1.**
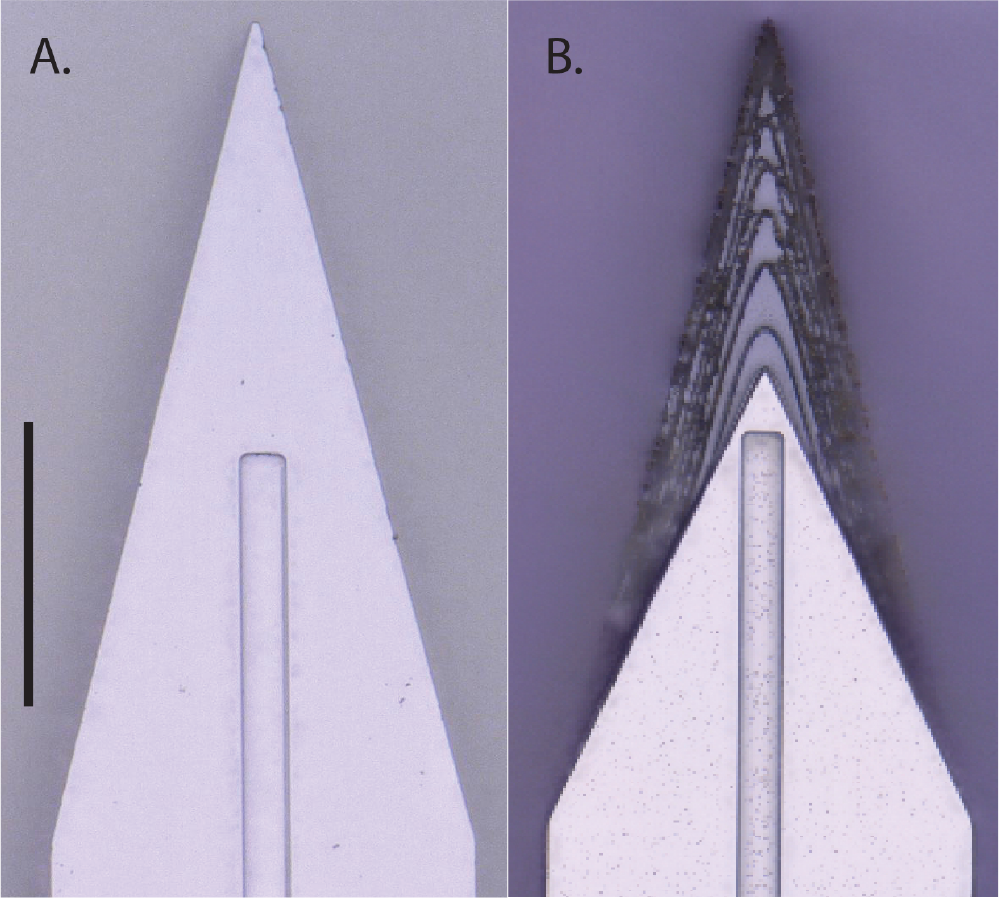
Top-down view by light microscope (scale bar = 50 µm) of (A) planar and (B) sharpened profiles on 2d- and 3d-sharpened silicon shuttles, respectively, oriented with tips toward the top of the figure, and wicking channel for PEG at midline.

**Figure 2.**
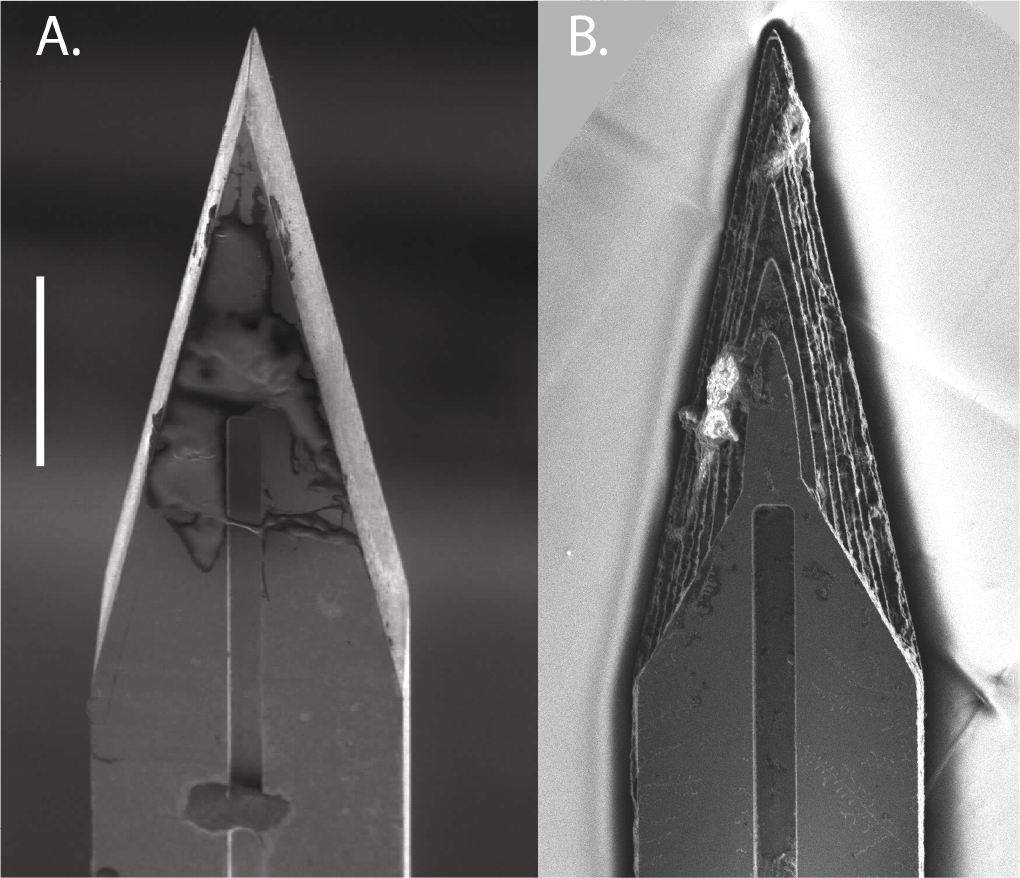
Top-down view by SEM for comparison of planar (A) and sharpened (B) shuttles (scale bar = 50 µm). Shuttles are oriented with tips toward the top of the figure and wicking channel for PEG at midline. Residue in (B) from previous insertion through neural tissue.

**Figure 3.**
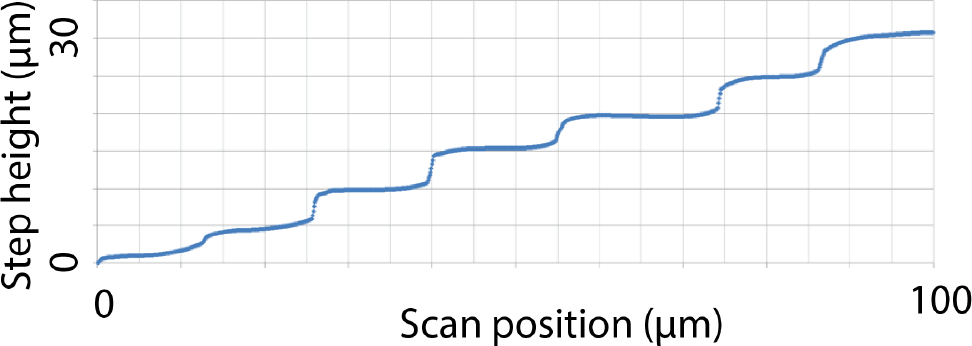
Profilometer scan of sharpened shuttle tip.

**Figure 4.**
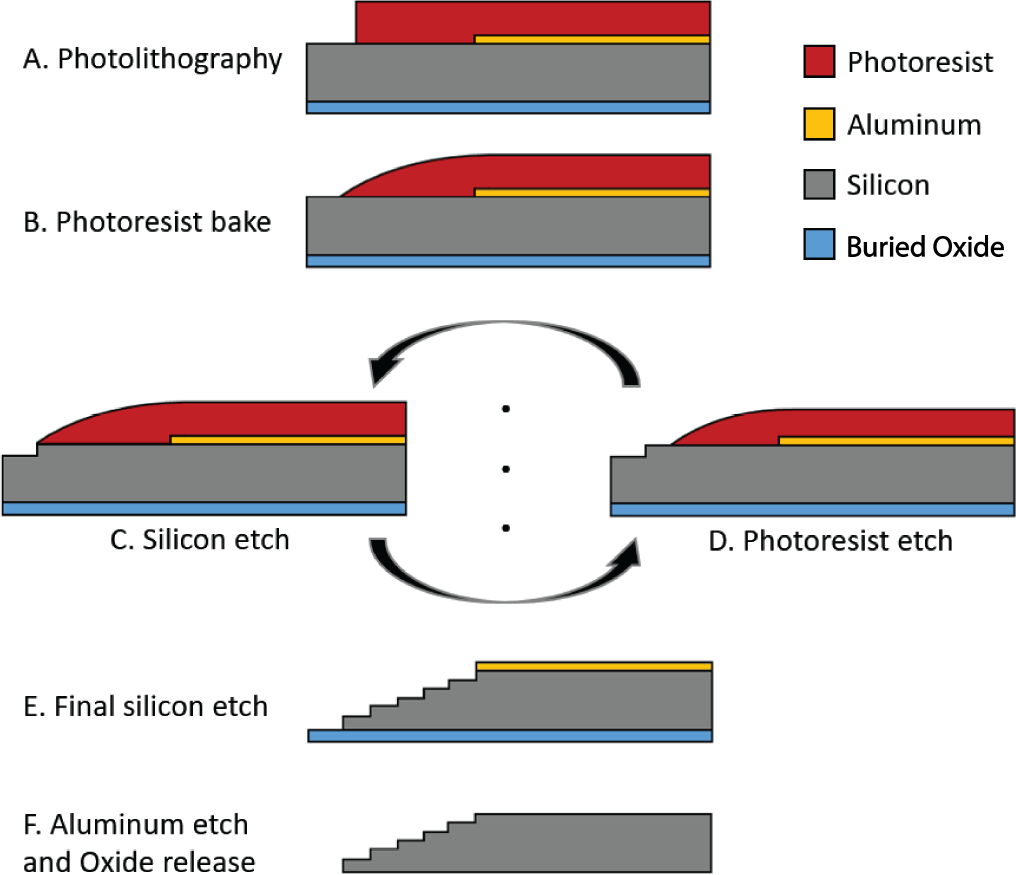
Process flow detail.

### 1.2 Polymer probe microfabrication

The probes chronically implanted in NAc and OFC were 32-channel, 2-shank devices, with 16 channels located at the tip of each 8 mm (NAc) or 6mm (OFC) shank, and design specifications as previously published (Chung et al., 2019).

### 1.3 Surgical insertion in vivo

For the comparison of force insertion measurements between planar and sharpened shuttles *in vivo*, two animals were used, ages 12 and 16 months. For sharpened shuttle-guided insertion of probes for chronic neural recordings, one animal, age 10 months, was used.

In all cases, animals were anesthetized with inhaled isoflurane plus a mixture of ketamine, xylazine, and atropine and were confirmed unresponsive to a foot pinch. The head was shaved and the animal transferred to the sterile surgery environment and head-fixed in a stereotactic frame. Body temperature was maintained throughout surgery by an isothermal pad beneath the animal. Following sterilization of the incision site and application of lidocaine as a local anaesthetic, a minimal incision was made through all skin layers at the top of the skull. The skin flaps were retracted and the tissues detached from the bone, preserving the attachment of the temporalis muscle to the temporal ridge. For the chronic recording animal, a dental drill was used to make 12 sub-penetrating skull holes along the temporal ridge. Into each of these holes, 0-80 titanium set screws (United Titanium, OH) were partially inserted, serving to anchor the implant to the skull. These were drilled with a rotating dental drill with carbide bur (SS White carbide bur, FG 2) at 6000 rpm, at an intermittent rate, with the drill removed completely from contact with the bone between bouts of drilling. Coordinates for target insertion sites were located using bregma and stereotactic coordinates. One craniotomy of approximately 2-3 mm diameter was made to expose the meninges over each of NAc (1.5 AP, ± 1.3 ML, mm) and OFC (+3.6 AP, ± 3.4 ML, mm) bilaterally (for chronic recordings), or each of hippocampus (−3.6 AP, ± 2.5 ML, mm) and OFC bilaterally (for force insertion measurements). The craniotomy technique was the same as for skull screw holes with the exception that the final ~500 µm of bone was removed using a smaller carbide bur (SS White carbide bur, FG ¼). The dura mater was maintained fully intact. Following *in vivo* data collection, animals were sacrificed using an overdose of pentobarbital sodium and phenytoin sodium (Euthasol, Virbac AH, Inc.).

### 1.4 In vivo insertion force measurements

Insertion force tests of shuttles without attached devices were performed over hippocampus and OFC bilaterally in each animal. In both experiments, insertion force measurements were performed with a precision load cell (FUTEK FSH02534) and voltage was read out through the digitizer (FUTEK FSH03633) to a computer at 100 samples/second. Rigid adapters between the motor, load cell, and insertion shuttle were 3D-printed with hard plastic from custom designs (PolyJetHD Blue, Stratasys Ltd.; Fig. 5). Insertions were conducted at a constant velocity of 50 µm/s using a micropositioner (Kopf model #2662).

**Figure 5.**
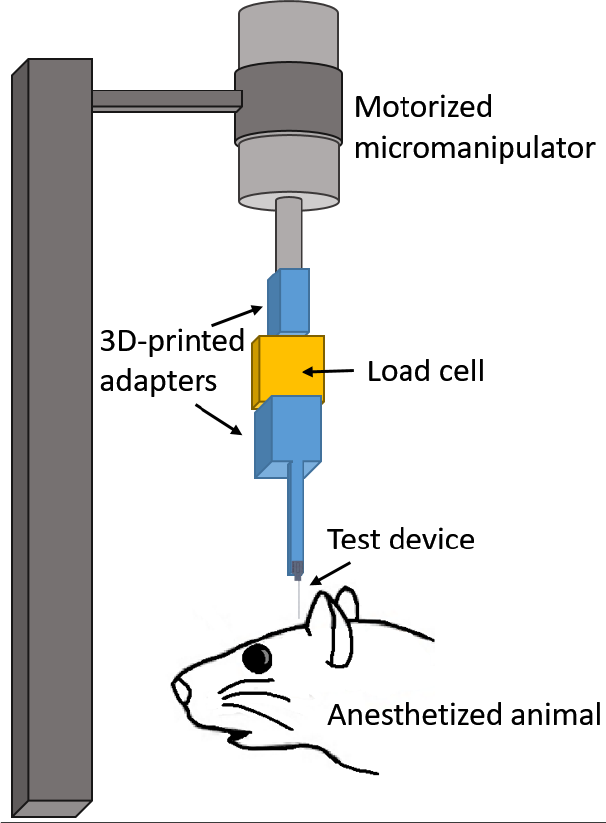
Test insertion apparatus for in vivo insertion force measurements. The test device was a single shank, 6mm long.

To match our typical surgical conditions, we tested serial insertions through dura in the same craniotomy, with at least 300 µm separation between insertion sites. Each insertion was performed with a shuttle through intact dura. The dura and brain were hydrated for the duration of the surgeries using hand irrigation with saline.

Shuttle insertions were classified as buckled if both (i) the force measured during insertion stopped increasing with time, despite the stepper motor continuing to advance the shuttle without dural penetration; and (ii) the shuttle visibly failed to enter the brain. A few shuttles exhibited buckling-like behavior during insertion, meeting criterion (i) above but not criterion (ii), as they proceeded to successfully penetrate the dura (Fig. 6). This occurred for one planar shuttle inserted through dura over right hippocampus, for one sharpened shuttle inserted over left OFC, and for one sharpened shuttle inserted over right OFC. The degree to which these devices buckled was judged by the experimenter to likely be sufficient to cause separation from an attached polymer array (maximum horizontal separation between probe and shuttle of ~50 µm or more), and these insertions were excluded from calculations of average maximum insertion force and brain compression.

**Figure 6.**
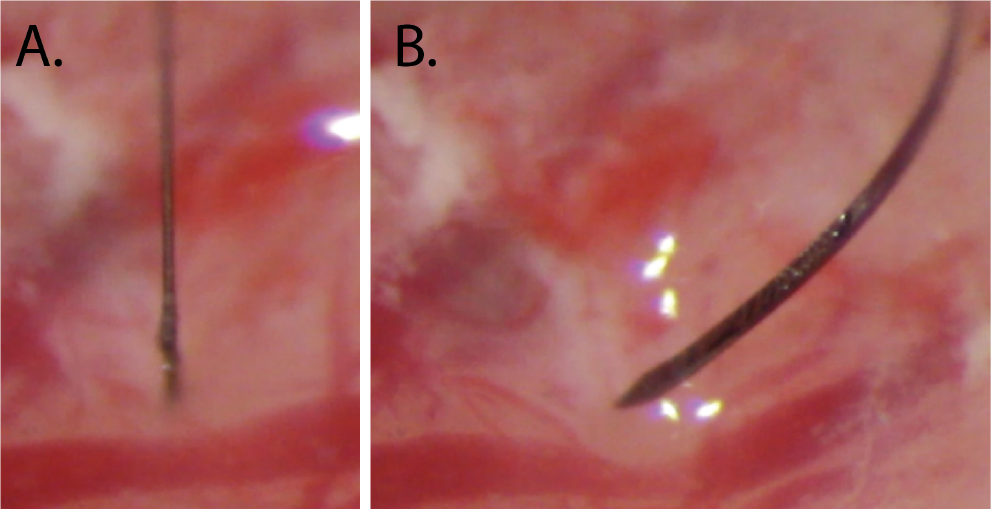
Example of a successful (A) and buckled (B) insertion attempt for sharpened silicon shuttles, 6mm.

Maximum insertion force was measured at the time of penetration, which was taken as the first absolute maximum of the force curve that preceded the first approximately infinite-slope curve characteristic of membrane penetration (Sridharan et al., 2013, Obaid et al., 2018). In a few cases, a slightly higher force was measured after initial penetration. In these cases we did not use the later timepoint to determine the maximum insertion force because, by then, the device had already begun to penetrate tissue and the total force measured thus included the frictional force as well.

Brain deflection distances were calculated by multiplying the insertion speed (50 µm/s) by the time (s) between dural contact and penetration. We estimated the time of dural contact by first calculating the mean and standard deviation of the (150-sample minimum) baseline period, during which the shuttle was still above tissue and contacting only air; we then determined the point at which a 5-sample sliding average of the measured force was at least one standard deviation above the baseline mean. The load cell compresses as a function of force applied, and the amount of compression was calibrated under a microscope and subtracted from deflection values calculated from force measurement data (resulting in a correction of 5 to 12 µm).

### 1.5 Imaging

Shuttles were imaged using Keyence Model VH-Z250R at ~1000X, with illuminated lighting on auto detect for white balance and brightness. Eight planar and eight sharpened shuttles were each imaged both before and after *in vivo* insertions through dura to verify that no shuttle breakage occurred in the brain during insertion or retraction. SEM images were taken of devices mounted on double-sided copper at magnification 300× with accelerating voltage 3kV using an APREO S Low Vacuum EDX Benchtop SEM (Fig. 2 a) or at magnification 650× with a Hitachi S-800 SEM (Fig. 2 b). The profilometer scan was taken using a Veeco Dektak profilometer.

### 1.6 Chronic recordings

All procedures were in accordance with guidelines from the University of California San Francisco Institutional Animal Care and Use Committee and US National Institutes of Health. For the tests of shuttles used with chronically implanted probes, the probe and shuttle were adhered to each other by PEG on the side opposite the recording contacts, as previously described for planar silicon shuttles. The meninges were hydrated between craniotomy completion and the time of insertion using hand irrigation with a saline drip. When all craniotomies were complete, a custom designed, 3D-printed plastic base piece was secured to the skull using dental acrylic attached to the 128-channel headstage (SpikeGadgets, LLC). These insertions were performed manually, lowering with a stereotax at approximately 50 µm/s until the probe penetrated the dura mater. Once in the brain, the device was lowered using a micromanipulator (MO-10, Narshige) at 10 µm/s until it was 1 mm above target depth, then at 5 µm/s until it was 500 µm above target depth, and at 3 µm/s until it reached target depth.

The four implant targets were: OFC bilaterally, at the same coordinates for the insertion force measurements, at depth - 4mm from brain surface; and NAc at depth −7 mm from brain surface. Following insertion, probes were secured to the base piece and the electronics connected. Once the probe reached target depth, polyimide “wings” glued to the probe perpendicular to its length were adhered to the base piece with acrylic, securing the probe at the target location. The shuttle was then detached from the probe by filling the base piece with saline, which dissolved the PEG at the probe-shuttle interface, thus allowing retraction of the shuttle without disrupting the probe. After all the probes were inserted, the base piece was drained of saline and filled with silicon elastomer (Dow Corning 3-4680) to prevent dural regrowth and scarring, brain swelling, and mechanical perturbation of the probes (Jackson and Muthuswamy, 2008). Kwik-sil (World Precision Instruments, LLC), dental acrylic, and a plastic case made of moldable plastic (ThermoMorph) was used to encase the devices. Post-surgically, meloxicam (Eloxiject, Henry Schein) and buprenorphine (Baytril, Reckitt Benckiser Healthcare) were given for pain, and a single dose of enrofloxacin (Baytril, Bayer Corporation) was given to prevent infection. Neural recordings were conducted in an approximately 1 square foot sleep box constructed of anti-static plastic and located in a recording room. Data were collected for 30 minutes or longer per day using the SpikeGadgets recording system (Trodes version 1.74), as previously described (Chung et al., 2019). The recording session analysed here was 1 hour long. Typically, the animal was asleep or quietly immobile during the recording period. The animal also ran a spatial navigation behaviour in epochs distinct from the sleep epoch analysed here. Recording quality was analysed at 95 days, as we have found previously that cell count using these polymer probes stabilizes by this time (Chung et al., 2019).

### 1.7 Neural data analysis

Data pre-processing was performed using custom Python and Matlab scripts. Common-average referencing was applied across the sixteen channels of each shank. Spike sorting was performed using the MountainSort software package (Chung et al., 2018, Chung et al., 2019), version 4.0. An initial round of automated sorting was performed with the following sorting parameters: detect sign =-1, detect threshold=3, clip size=100, adjacency radius=200 µm. The raw data were filtered between 600 and 6000 Hz. The detect interval was set to 10 samples and the first 10 principal components were used. For units identified using these parameters, only those with cluster quality metrics above the following thresholds were included as single units: firing rate threshold=0.01 Hz, isolation threshold=0.96, noise overlap threshold=0.03, peak signal-to-noise ratio threshold=1.5. The rest were marked as multi-unit activity. All identified units, including those marked as multi-unit activity, were manually inspected and curated: in MountainView software, clusters that did not appear to be single units based on refractory period violations (*i.e.*, frequent spiking within 2 ms of the last spike) were rejected; multiple clusters that were identified as corresponding to the same unit based on a combination of firing rates, waveforms, peak channels, and temporal cross-correlograms were merged, and all cluster metrics listed above were re-calculated. Clusters that passed the curation metric thresholds were accepted.

## Results

### 1.1 Planar and sharpened shuttle insertion forces

We first tested single-shank shuttles with either a flat profile (planar; N = 5) or a sharpened profile (sharpened; N = 8) but otherwise identical dimensions (6mm × 30µm × 80µm, Fig. 1 and Fig. 2) in transdural insertion to the hippocampus (Fig. 7 a). Shuttles were lowered by micropositioner at a constant speed of 50 µm/s and insertion force was measured as the shuttle contacted dura and either inserted or buckled (Fig. 6).

**Figure 7.**
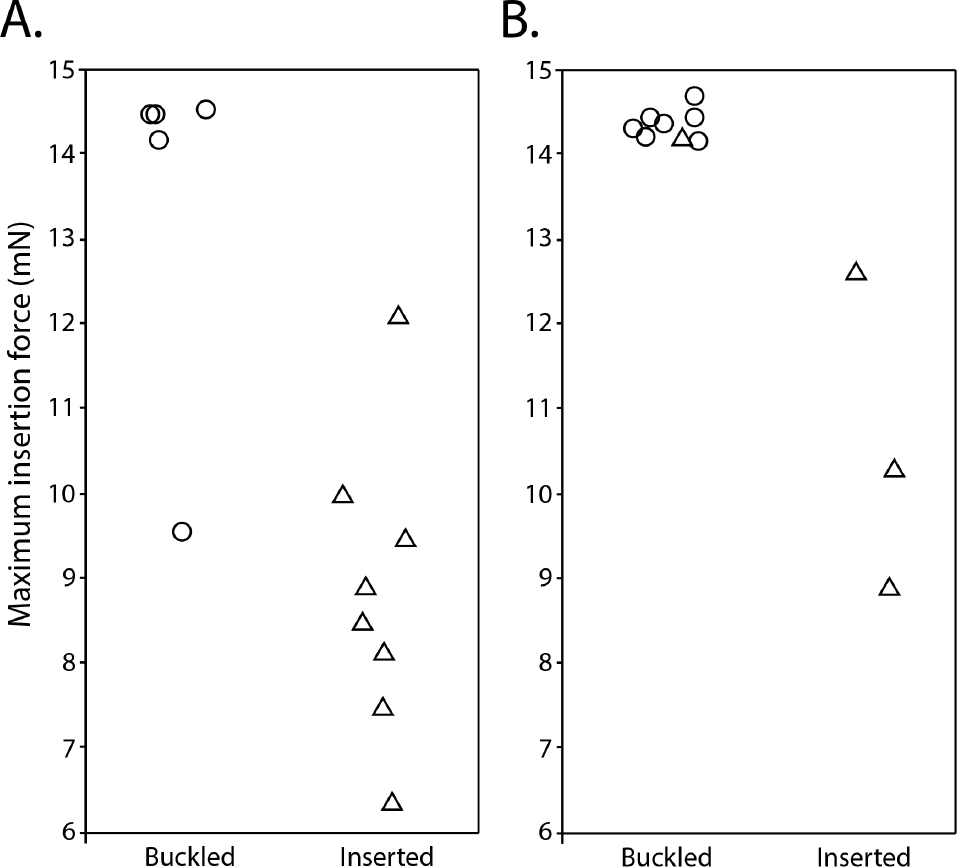
Maximum insertion force for sharpened (triangles) and planar (circles) shuttles inserted to (A) hippocampus (B) OFC. Y-axis=maximum insertion force in mN. Random jitter applied along x-axis for visualization.

Sharpened shuttles penetrated dura (8/8, 100%) but planar shuttles did not (0/5, 0%). One planar shuttle successfully penetrated dura over the right hippocampus, but was excluded from subsequent calculations because it penetrated dura only after buckling to an extent that would likely have caused separation from an attached polymer probe. Fig. 7 a shows the maximum insertion force for each insertion test over hippocampus.

For successfully inserted sharpened shuttles, the average insertion force was 8.86 ± 0.61 mN (N=8, ± s.e.m.; Fig. 7 a, right). The insertion force traces for shuttles that successfully penetrated the dura showed increasing force, followed by a characteristic series of rapid decrements between local maxima, in a sawtooth pattern (Fig. 8). In a few cases, a slightly higher force was measured following initial penetration (Fig. 8, panels 3, 9, 10, 11 from top left to bottom right), likely due to increased friction force after the device had begun to penetrate tissue. In these cases we took the first local maximum as the maximum insertion force. In contrast, failed penetrations with unsharpened shuttles resulted in buckling and had an average buckling force of 13.42 ± 0.97 mN (N = 5, ± s.e.m.; Fig. 7 a, left).

**Figure 8.**
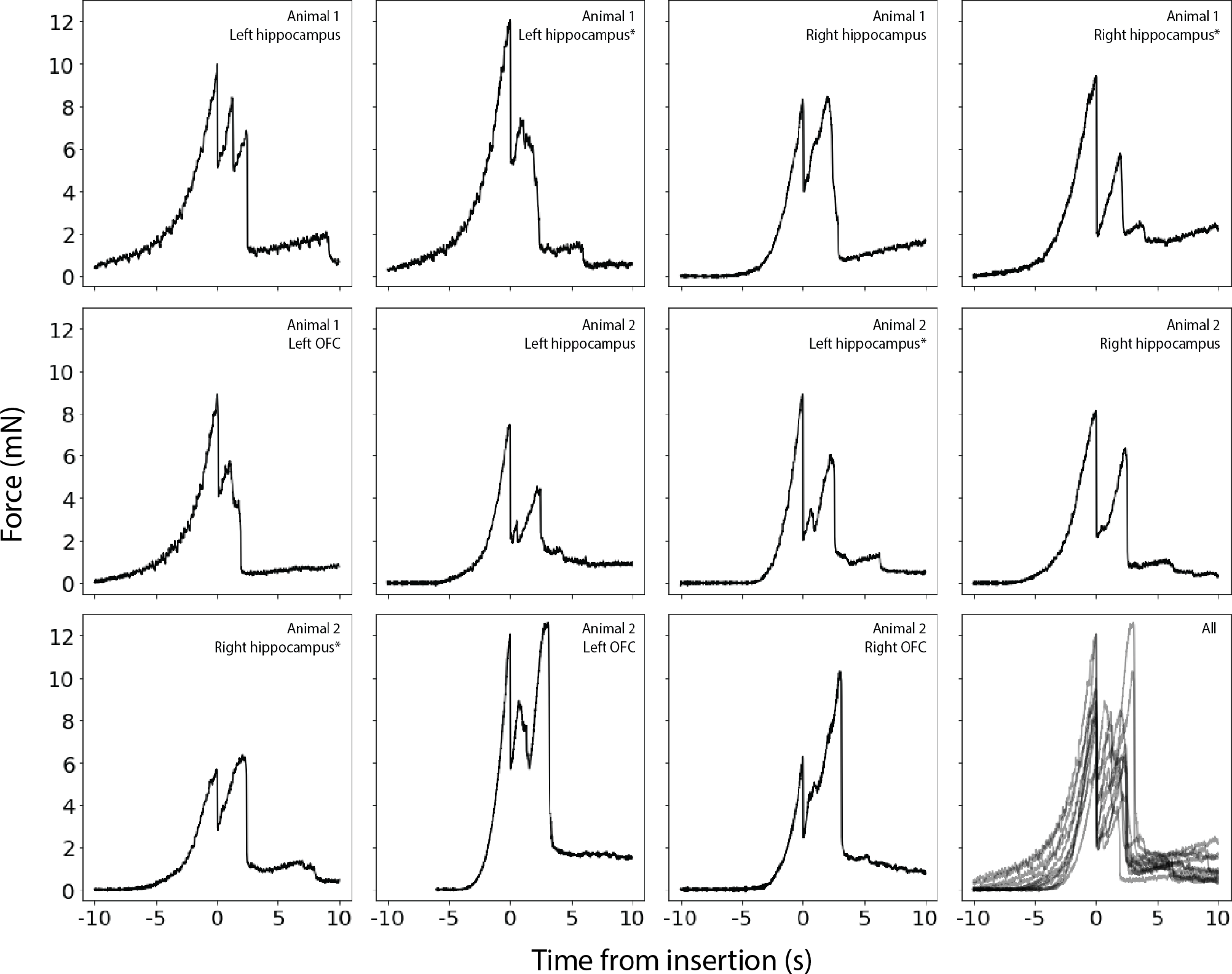
Raw force traces for all sharpened shuttles inserted through dura. X-axes in seconds, with traces aligned to 0 as the timepoint of maximum force, y-axes in mN. The animal and area of insertion are labelled in the upper right of each panel; plot at lower right shows all insertion force measurements depicted here overlaid. If the insertion was through a different area of a craniotomy where a section of dura at least 300 µm away was already used for a test insertion, the insertion site is marked with an asterisk.

Our hippocampal target was located approximately halfway between bregma and lambda, and halfway between the midline and the temporal ridge. As a test of the effectiveness of our sharpened shuttles through thicker dura, we next performed the same tests over the OFC, where the dura is thicker and more difficult to penetrate. Tests were performed for OFC as for hippocampus (Fig. 7 b). Sharpened profile shuttles penetrated dura 3/4 times (75%) while planar shuttles penetrated dura 0/7 times (0%). This slightly worse performance over OFC than hippocampus is expected for tougher dura. Two sharpened shuttle insertions, one over left OFC and one over right OFC, successfully penetrated the brain only after buckling to an extent that would likely have caused separation from an attached device and were excluded from subsequent calculations. Figure 7 b shows the maximum insertion force for each insertion test over OFC.

For successfully inserted sharpened shuttles over OFC, the average insertion force was 10.61 ± 1.08 mN (N=3, ± s.e.m.; Fig. 7 b, right; insertion profiles also included in Fig. 8), which is not significantly different than for hippocampus (Welch’s t-test: p=0.24). In contrast, failed penetrations (buckled) with unsharpened shuttles resulted in buckling and had an average buckling force of 14.35 ± 0.07 mN (N = 7, ± s.e.m.; Fig. 7 b, left).

For successfully inserted sharpened shuttles over OFC, the average insertion force was 10.61 ± 1.08 mN (N=3, ± s.e.m.; Fig. 7 b, right; insertion profiles also included in Fig. 8), which is not significantly different than for hippocampus (Welch’s t-test: p=0.24). In contrast, failed penetrations (buckled) with unsharpened shuttles resulted in buckling and had an average buckling force of 14.35 ± 0.07 mN (N = 7, ± s.e.m.; Fig. 7 b, left).

### 1.2 Brain compression on transdural shuttle insertion

We calculated the degree of brain compression for sharpened shuttles successfully inserted through dura (*i.e.*, those insertions included in Fig. 8). On average, the calculated compression for sharpened shuttle insertions through dura was 389.9 ± 159.6 µm (N=8, ± s.e.m.) for hippocampus and 302.5 ± 158.5 µm (N=3, ± s.e.m.) for OFC.

### 1.3 Imaging before and after insertion

We imaged 8 sharpened and 8 planar shuttles before and after *in vivo* insertion. We observed no breakage or other discernible damage to any of the shuttle tips, indicating that sharpened shuttles, despite having a tip diameter of less than 3µm in width and thickness, maintain their structural integrity and do not break off in the brain (Fig. 9).

**Figure 9.**
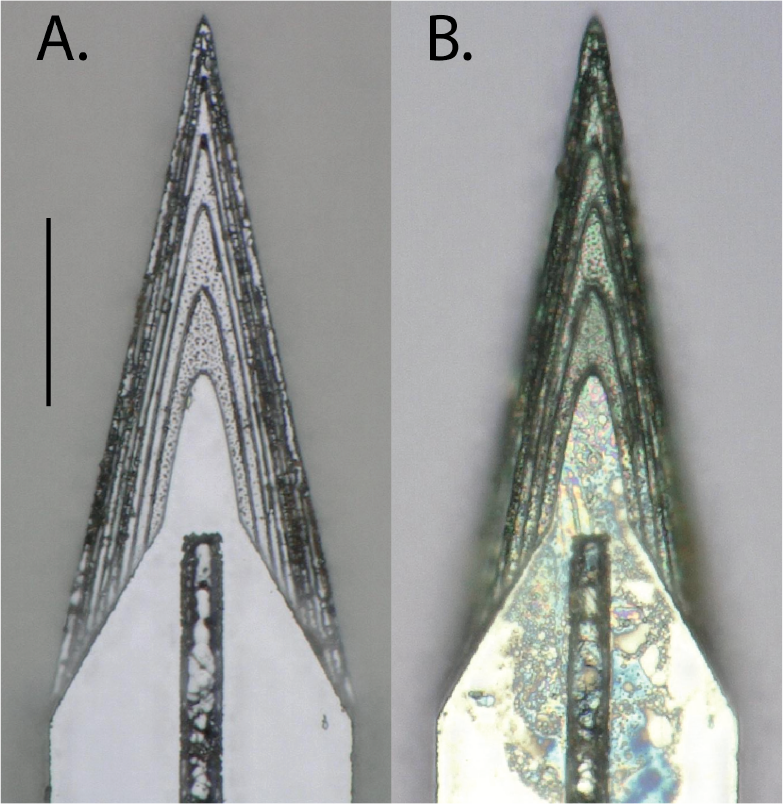
Top-down view of the same sharpened shuttle. A. Before insertion through dura. B. After insertion. Scale bar=50 µm.

### 1.4 Neural recordings using sharpened shuttles

During insertion of recording devices adhered to sharpened shuttles, devices entered the brain with minimal compression of the dura and brain. After dural penetration using a stereotax, the device was inserted to its final depth using a micromanipulator.

We observed that single units were detected on all six implanted shanks (16 channels each) for at least 90 days. We selected a single epoch on day 95 post-implant to perform spike sorting on one of the shanks targeted to the left OFC. This shank was selected for its electrode quality prior to insertion (no shorts or dead channels) and relatively low noise levels (Fig. 10 a). We identified 18 single units on this 16-channel shank, for an average of approximately one unit per electrode (Fig. 10 b). This is similar to what we have previously observed for high-quality multi-electrode arrays using planar shuttles (Chung et al., 2019). Single units that passed curation standards were evenly distributed over the recording channels, with no apparent systematic difference in quality, from the top to bottom of the shank.

**Figure 10.**
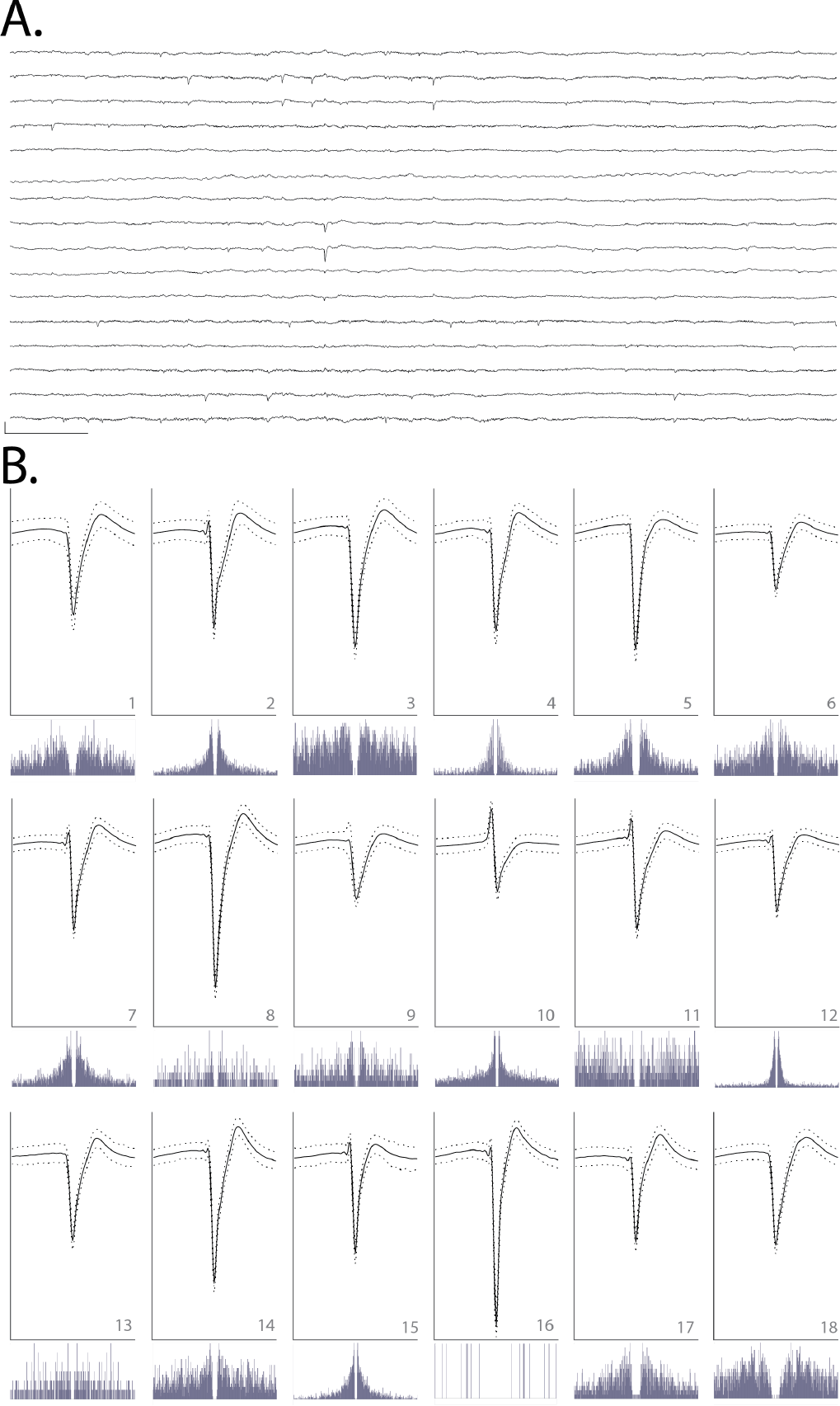
**A**. Raw, 150-millisecond local field potential recorded on each of the 16 channels of one shank targeted to OFC, 95 days after implant. Channels 1-16 are ordered from top to bottom. Channels 1-8 correspond to electrodes on the left side of the probe shank, from top to bottom; channels 9-16 correspond to electrodes on the right side of the device, from bottom to top (i.e., channels 1 and 16 are at similar depth). Scale bar, bottom left = 1 mV vertical, 15 milliseconds horizontal. **B**. Each of 18 units detected on the 16-channel shank shown in A. Odd rows: average waveforms (solid line) +/− one standard deviation (dashed line) for units 1-18, numbered in bottom right of each panel. For each waveform, vertical scale bar = 2.5 millivolts, horizontal scale bar = 100 samples (at 30 kHz), or approximately 3.33 milliseconds. Even rows: spike auto-correlograms for the unit shown above, spanning 100 milliseconds, in 0.5-millisecond bins.

## Discussion

Previous work has demonstrated the benefits of sharpened device tips in reducing insertion force (Sharp et al., 2009, Fekete et al., 2015, Obaid et al., 2018) and penetrating dura (Hosseini et al., 2007, Fekete et al., 2015, Sharp et al., 2009). Here, we sharpen not only in two dimensions but three, resulting in a more gradually increasing cross-section (Chen et al., 2017, Fekete et al., 2015). This is, to our knowledge, the first demonstration of a dural-penetrating, polymer device insertion method to yield chronic single unit recordings. It also allowed us to directly compare shuttles made of the same material (silicon) that were sharp in three versus two dimensions.

We have shown that sharpened silicon shuttles result in low insertion forces and brain compression. The expected buckling force, also known as the critical load, of our silicon shuttle can be calculated with Euler’s critical load equation:

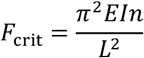

where *E* is the Young’s modulus of silicon, *I* is the moment of inertia for the cross-sectional area of the shuttle, *n* is a factor accounting for end conditions, and *L* is the length of the shuttle. The moment of inertia *I* of a rectangular cross-section is *ab*^3^/12 where a and b are the longer and shorter side lengths, respectively. For translation fixed, rotation fixed and translation fixed, rotation free (brain side) end conditions, *n*=2 and the expected critical load

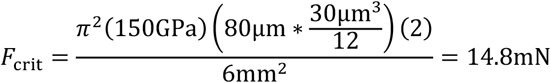

is very close to our measured result of 13.42 mN for hippocampus and 14.35 mN for OFC. Due to the deformable nature of the brain, the boundary condition on the brain side is not completely translation fixed, with true *n* less than 2 and true F_crit_ expected to be slightly lower than calculated.

Our maximum insertion force for transdural insertion of sharpened shuttles of aproximately 10 mN is lower than previous reports for planar devices (*e.g.*, 41 mN with silicon probes; Hosseini et al., 2007) and similar to the 11 ± 2 mN reported previously in a proof-of-concept study in which a less flexible, silicon-specific etch technique was used to insert but not record from silicon probes (Fekete et al., 2015). For comparison, previously reported peak insertion forces for rat pia-only insertions are approximately 5 mN (Paralikar and Clement*, 2008, Fekete et al., 2015).

Recently, an insertion method using a diamond shuttle tested rat transdural insertion at a variety of speeds, finding a minimum average compression of 488 µm at insertion speed 10 µm/sec with 200 Hz vibration (Na et al., 2018). At approximately 300-400 µm of compression without piezovibration, our method is comparable. To our knowledge, these are the lowest brain compression values for a working device through dura. It is a reasonable expectation that for surgeries where performing a durotomy is preferable, these devices can be implanted into the brain through pia with even lower brain compression. This use could be particularly valuable for superficial cortical sites in which recording quality is easily compromised by compression of the brain (Rennaker et al., 2005).

Force and tissue compression are important metrics to gauge neural tissue damage, but a single unit count weeks after implant is required to fully demonstrate a neural recording technology for chronic use. Recently, promising strategies for insertion of flexible polymer devices through rat dura have been demonstrated (Shoffstall et al., 2018, Zhao et al., 2019), but such methods have not yet reported chronic, single-unit recordings in freely behaving animals. Here, we evaluated our single unit count at 95 days. At that timepoint we were able to record high quality single units on 11 of 16 (69%) channels, with a total of 18 sorted units on 16 channels, or an average of 1 high-quality unit per electrode. In comparison, the other recent technology that reports a single unit count is the dural penetrating diamond shuttle for delivery of a flexible array, which yielded acute neural recordings of 20 units over 60 channels, or an average of 0.33 units per electrode (Na et al., 2018). Another exciting alternative to manual durotomy is laser microablation, which has been used preceding robotic insertion of thin-film polymer probes. The longest recordings reported using this method were taken at two months post-implant, with approximately 40 percent of channels recording single-unit action potentials (Hanson et al., 2019).

Further reduction in brain compression may be achieved by fabrication of even sharper profile tips (Obaid et al., 2018), which our fabrication method can achieve with minimal adjustment. Another way to reduce brain compression could be to change insertion speed. A range of speeds, including meters-per-second scale pneumatic insertion (Black et al., 2018), have yielded dural penetration (Fekete et al., 2015) and neural recordings (Zhao et al., 2019, Hosseini et al., 2007, P. Rousche, 1992), with the optimal speed dependent on tip shape and features in the target tissue, such as vessels (Bjornsson et al., 2006). Retraction speed, too, has been varied; for example, the ballistic retraction method used in combination with needle-and-thread insertion to prevent displacement of electrodes from target depth has, as previously discussed, yielded chronic neural recordings (Hanson et al., 2019). In addition to determining the optimal insertion and retraction speeds, incorporating fast axioaxial vibration with relatively slow insertion speed could also aid in insertion with minimal brain compression (Na et al., 2018). Additionally, application of collagenase to the dura could make it more easily penetrable (Paralikar and Clement*, 2008), though this process is slow and titration to determine an appropriate concentration presents a challenge, as the dural membrane thickness varies with target location and age (Fekete et al., 2015).

Another complementary insertion strategy uses an insertion guide, as recently demonstrated for insertion of SMP devices through rat dura (Shoffstall et al., 2018). Inspired by the labium that guides the proboscis of the female mosquito, a guide at the insertion site increases the critical buckling load of the inserted device itself. Such a strategy could be used in combination with the sharpened shuttles validated here.

Although some stiff devices can be inserted directly through the meninges (P. Rousche, 1992, Escamilla-Mackert et al., 2009), the damage to the blood brain barrier seen with any implanted device can be severe, and compression of brain tissue can have an effect similar to traumatic brain injury (Kozai et al., 2015). It is also possible that the design and fabrication technique presented here could be used for silicon electrode arrays themselves, either for dural penetration or to reduce brain compression in clinical and non-human primate implants.

## Conclusion

Recording more neurons in distributed circuits will likely lead to scientific insights that are unachievable with a smaller number of neurons (Stevenson and Kording, 2011, Buzsaki, 2004). Polymer probes are among the most promising methods for chronic, large-scale neural recordings, but their insertion through the tough protective membranes of the central nervous system is challenging and currently limits their broad use and effectiveness. Here, we have validated for chronic recordings the first dural-penetrating shuttle in combination with a modular polymer probe-based recording platform. This method shows limited brain compression and obviates the need for a durotomy in rats and other model organisms with similar dural tensile strength. Maintaining intact dura will reduce post-surgical edema, likely increasing accuracy in depth-targeting of the electrode arrays. This is critical for fixed, non-drivable arrays, particularly for targets with a small dorsoventral extent. The number and quality of single units we have recorded with this system is comparable to what we have previously recorded in the OFC using planar shuttles, inserted through pia only (Chung et al., 2018). It is our hope that this method to more efficiently implant polymer devices for high-density, chronic neural recordings will enable experimentalists to address compelling open questions in neuroscience.

## Acknowledgements

We thank Thomas J. Davidson and Anna K. Gillespie for useful discussion regarding spike sorting, Clay Smyth for force measurement and imaging assistance, and Anna Kiseleva for assistance in animal care. This work was supported by National Institute of Mental Health (NIMH) award number F30MH115582 (H.R.J.), NIMH award number F30MH109292 (J.E.C.), NIMH award number F31MH112335 (D.K.R.), National Institute of General Medical Sciences Medical Scientist Training Program grant #T32GM007618 (H.R.J., J.E.C.), NIMH grants U01 NS090537 (LLNL and L.M.F.) and UF1 NS107667 (L.M.F.) and the Howard Hughes Medical Institute (L.M.F.). This work was performed under the auspices of the U.S. Department of Energy by Lawrence Livermore National Laboratory under contract DE-AC52-07NA27344. LLNL-JRNL-770828.

## References

Bjornsson, C. S., Oh, S. J., Al-Kofahi, Y. A., Lim, Y. J., Smith, K. L., Turner, J. N., De, S., Roysam, B., Shain, W. & Kim, S. J. 2006. Effects of insertion conditions on tissue strain and vascular damage during neuroprosthetic device insertion. J Neural Eng, 3, 196–207.

Black, B. J., Kanneganti, A., Joshi-Imre, A., Rihani, R., Chakraborty, B., Abbott, J., Pancrazio, J. J. & Cogan, S. F. 2018. Chronic recording and electrochemical performance of Utah microelectrode arrays implanted in rat motor cortex. J Neurophysiol, 120, 2083–2090.

Buzsaki, G. 2004. Large-scale recording of neuronal ensembles. Nat.Neurosci., 7, 446–451.

Chen, S., Fan, J. L., Chung, J. E., Joo, H. R., Pebbles, J., Frank, L. M. & Tolosa, V. 2017. 3D-Sharpened, Microfabricated Tool for Insertion of Flexible Electrode Arrays into Brain. 39th Annual International Conference of the IEEE Engineering in Medicine and Biology Society. Jeju Island, Korea.

Chung, J. E., Joo, H. R., Fan, J. L., Liu, D. F., Barnett, A. H., Chen, S., Geaghan-Breiner, C., Karlsson, M. P., Karlsson, M., Lee, K. Y., Liang, H., Magland, J. F., Pebbles, J. A., Tooker, A. C., Greengard, L. F., Tolosa, V. M. & Frank, L. M. 2019. High-Density, Long-Lasting, and Multi-region Electrophysiological Recordings Using Polymer Electrode Arrays. Neuron, 101, 21–31.e5.

Chung, J. E., Magland, J. F., Barnett, A. H., Tolosa, V. M., Tooker, A. C., Lee, K. Y., Shah, K. G., Felix, S. H., Frank, L. M. & Greengard, L. F. 2017. A Fully Automated Approach to Spike Sorting. Neuron, 95, 1381–1394 e6.

D. Kipke, R. V., J. Williams, J. Hetke 2003. Silicon-Substrate Intracortical Microelectrode Arrays for Long-Term Recording of Neuronal Spike Activity in Cerebral Cortex. IEEE TRANSACTIONS ON NEURAL SYSTEMS AND REHABILITATION ENGINEERING, 11, 151–155.

Dixon, C. E., Clifton, G. L., Lighthall, J. W., Yaghmai, A. A. & Hayes, R. L. 1991. A controlled cortical impact model of traumatic brain injury in the rat. J Neurosci Methods, 39, 253–62.

Edell, D. J., Toi, V. V., Mcneil, V. M. & Clark, L. D. 1992. Factors influencing the biocompatibility of insertable silicon microshafts in cerebral cortex. IEEE Trans Biomed Eng, 39, 635–43.

Escamilla-Mackert, T., Langhals, N. B., Kozai, T. D. & Kipke, D. R. 2009. Insertion of a three dimensional silicon microelectrode assembly through a thick meningeal membrane. Conf Proc IEEE Eng Med Biol Soc, 2009, 1616–8.

Fekete, Z., Nemeth, A., Marton, G., Ulbert, I. & Pongracz, A. 2015. Experimental study on the mechanical interaction between silicon neural microprobes and rat dura mater during insertion. J Mater Sci Mater Med, 26, 70.

Felix, S., Shah, K., George, D., Tolosa, V., Tooker, A., Sheth, H., Delima, T. & Pannu, S. 2012. Removable silicon insertion stiffeners for neural probes using polyethylene glycol as a biodissolvable adhesive. Conf Proc IEEE Eng Med Biol Soc, 2012, 871–4.

Fu, T. M., Hong, G. S., Zhou, T., Schuhmann, T. G., Viveros, R. D. & Lieber, C. M. 2016. Stable long-term chronic brain mapping at the single-neuron level. Nature Methods, 13, 875-+.

Hanson, T. L., Diaz-Botia, C. A., Kharazia, V., Maharbiz, M. M. & Sabes, P. N. 2019. The “sewing machine” for minimally invasive neural recording. bioRxiv, 578542.

Harris, J. P., Capadona, J. R., Miller, R. H., Healy, B. C., Shanmuganathan, K., Rowan, S. J., Weder, C. & Tyler, D. J. 2011. Mechanically adaptive intracortical implants improve the proximity of neuronal cell bodies. J Neural Eng, 8, 066011.

Hong, G. & Lieber, C. M. 2019. Novel electrode technologies for neural recordings. Nat Rev Neurosci.

Hosseini, N. H., Hoffmann, R., Kisban, S., Stieglitz, T., Paul, O. & Ruther, P. 2007. Comparative study on the insertion behavior of cerebral microprobes. Conf Proc IEEE Eng Med Biol Soc, 2007, 4711–4.

Jackson, N. & Muthuswamy, J. 2008. Artificial dural sealant that allows multiple penetrations of implantable brain probes. J Neurosci Methods, 171, 147–52.

Jensen, W., Yoshida, K. & Hofmann, U. G. 2006. In-vivo implant mechanics of Flexible, silicon-based ACREO microelectrode arrays in rat cerebral cortex. IEEE Trans Biomed Eng, 53, 934–40.

Jeong, J. W., Shin, G., Park, S. I., Yu, K. J., Xu, L. & Rogers, J. A. 2015. Soft materials in neuroengineering for hard problems in neuroscience. Neuron, 86, 175–86.

Jun, J. J., Steinmetz, N. A., Siegle, J. H., Denman, D. J., Bauza, M., Barbarits, B., Lee, A. K., Anastassiou, C. A., Andrei, A., Aydin, C., Barbic, M., Blanche, T. J., Bonin, V., Couto, J., Dutta, B., Gratiy, S. L., Gutnisky, D. A., Hausser, M., Karsh, B., Ledochowitsch, P., Lopez, C. M., Mitelut, C., Musa, S., Okun, M., Pachitariu, M., Putzeys, J., Rich, P. D., Rossant, C., Sun, W. L., Svoboda, K., Carandini, M., Harris, K. D., Koch, C., O’Keefe, J. & Harris, T. D. 2017. Fully integrated silicon probes for high-density recording of neural activity. Nature, 551, 232–236.

Khilwani, R., Gilgunn, P. J., Kozai, T. D., Ong, X. C., Korkmaz, E., Gunalan, P. K., Cui, X. T., Fedder, G. K. & Ozdoganlar, O. B. 2016. Ultra-miniature ultra-compliant neural probes with dissolvable delivery needles: Design, fabrication and characterization. Biomed Microdevices, 18, 97.

Kim, B. J., Kuo, J. T., Hara, S. A., Lee, C. D., Yu, L., Gutierrez, C. A., Hoang, T. Q., Pikov, V. & Meng, E. 2013. 3D Parylene sheath neural probe for chronic recordings. J Neural Eng, 10, 045002.

Kozai, T. D., Gugel, Z., Li, X., Gilgunn, P. J., Khilwani, R., Ozdoganlar, O. B., Fedder, G. K., Weber, D. J. & Cui, X. T. 2014. Chronic tissue response to carboxymethyl cellulose based dissolvable insertion needle for ultra-small neural probes. Biomaterials, 35, 9255–68.

Kozai, T. D., Jaquins-Gerstl, A. S., Vazquez, A. L., Michael, A. C. & Cui, X. T. 2015. Brain tissue responses to neural implants impact signal sensitivity and intervention strategies. ACS Chem Neurosci, 6, 48–67.

Kozai, T. D. & Kipke, D. R. 2009. Insertion shuttle with carboxyl terminated self-assembled monolayer coatings for implanting flexible polymer neural probes in the brain. J Neurosci Methods, 184, 199–205.

Kozai, T. D., Langhals, N. B., Patel, P. R., Deng, X., Zhang, H., Smith, K. L., Lahann, J., Kotov, N. A. & Kipke, D. R. 2012. Ultrasmall implantable composite microelectrodes with bioactive surfaces for chronic neural interfaces. Nat Mater, 11, 1065–73.

Lacour, S. P., Courtine, G. & Guck, J. 2016. Materials and technologies for soft implantable neuroprostheses. Nature Reviews Materials, 1.

Lecomte, A., Descamps, E. & Bergaud, C. 2018. A review on mechanical considerations for chronically-implanted neural probes. J Neural Eng, 15, 031001.

Lee, H. C., Ejserholm, F., Gaire, J., Currlin, S., Schouenborg, J., Wallman, L., Bengtsson, M., Park, K. & Otto, K. J. 2017. Histological evaluation of flexible neural implants; flexibility limit for reducing the tissue response? J Neural Eng, 14, 036026.

Liu, J., Fu, T. M., Cheng, Z., Hong, G., Zhou, T., Jin, L., Duvvuri, M., Jiang, Z., Kruskal, P., Xie, C., Suo, Z., Fang, Y. & Lieber, C. M. 2015. Syringe-injectable electronics. Nat Nanotechnol, 10, 629–636.

Lo, M. C., Wang, S., Singh, S., Damodaran, V. B., Kaplan, H. M., Kohn, J., Shreiber, D. I. & Zahn, J. D. 2015. Coating flexible probes with an ultra fast degrading polymer to aid in tissue insertion. Biomed Microdevices, 17, 34.

Lozano, A. M., Lipsman, N., Bergman, H., Brown, P., Chabardes, S., Chang, J. W., Matthews, K., Mcintyre, C. C., Schlaepfer, T. E., Schulder, M., Temel, Y., Volkmann, J. & Krauss, J. K. 2019. Deep brain stimulation: current challenges and future directions. Nat Rev Neurol.

Luan, L., Wei, X., Zhao, Z., Siegel, J. J., Potnis, O., Tuppen, C. A., Lin, S., Kazmi, S., Fowler, R. A., Holloway, S., Dunn, A. K., Chitwood, R. A. & Xie, C. 2017. Ultraflexible nanoelectronic probes form Reliable, glial scar-free neural integration. Sci Adv, 3, e1601966.

Luo, Y. H. & Da Cruz, L. 2016. The Argus((R)) II Retinal Prosthesis System. Prog Retin Eye Res, 50, 89–107.

Maikos, J. T., Elias, R. A. & Shreiber, D. I. 2008. Mechanical properties of dura mater from the rat brain and spinal cord. J Neurotrauma, 25, 38–51.

Mols, K., Musa, S., Nuttin, B., Lagae, L. & Bonin, V. 2017. In vivo characterization of the electrophysiological and astrocytic responses to a silicon neuroprobe implanted in the mouse neocortex. Sci Rep, 7, 15642.

Na, K., Sperry, Z. J., Lu, J., Voeroeslakos, M., Parizi, S. S., Bruns, T. M., Yoon, E. & Seymour, J. P. 2018. Novel diamond shuttle to deliver flexible bioelectronics with reduced tissue compression. bioRxiv, 435800.

Nicolelis, M. A., Dimitrov, D., Carmena, J. M., Crist, R., Lehew, G., Kralik, J. D. & Wise, S. P. 2003. Chronic, Multisite, multielectrode recordings in macaque monkeys. Proc Natl Acad Sci U S A, 100, 11041–11046.

Obaid, A. M., Wu, Y.-W., Hanna, M.-E., Nix, W. D., Ding, J. B. & Melosh, N. A. 2018. Ultra-sensitive measurement of brain penetration with microscale probes for brain machine interface considerations. bioRxiv, 454520.

P. Rousche, R. N. 1992. A method for pneumatically inserting an array of penetrating electrodes into cortical tissue. Annals of Biomedical Engineering, 20, 413–422.

Paralikar, K. J. & Clement, R. S.* 2008. Collagenase-Aided Intracortical Microelectrode Array Insertion: Effects on Insertion Force and Recording Performance. IEEE Transactions on Biomedical Engineering, 55, 2258–2267.

Patel, P. R., Na, K., Zhang, H., Kozai, T. D., Kotov, N. A., Yoon, E. & Chestek, C. A. 2015. Insertion of linear 8.4 µm diameter 16 channel carbon fiber electrode arrays for single unit recordings. J Neural Eng, 12, 046009.

Polikov, V. S., Tresco, P. A. & Reichert, W. M. 2005. Response of brain tissue to chronically implanted neural electrodes. J Neurosci Methods, 148, 1–18.

Rennaker, R. L., Street, S., Ruyle, A. M. & Sloan, A. M. 2005. A comparison of chronic multi-channel cortical implantation techniques: manual versus mechanical insertion. J Neurosci Methods, 142, 169–76.

Rousche, P. J. & Normann, R. A. 1998. Chronic recording capability of the Utah Intracortical Electrode Array in cat sensory cortex. J Neurosci Methods, 82, 1–15.

Schuhmann, T. G., Jr., Zhou, T., Hong, G., Lee, J. M., Fu, T. M., Park, H. G. & Lieber, C. M. 2018. Syringe-injectable Mesh Electronics for Stable Chronic Rodent Electrophysiology. J Vis Exp.

Seymour, J. P. & Kipke, D. R. 2007. Neural probe design for reduced tissue encapsulation in CNS. Biomaterials, 28, 3594–607.

Sharp, A. A., Ortega, A. M., Restrepo, D., Curran-Everett, D. & Gall, K. 2009. In vivo penetration mechanics and mechanical properties of mouse brain tissue at micrometer scales. IEEE Trans Biomed Eng, 56, 45–53.

Shoffstall, A. J., Srinivasan, S., Willis, M., Stiller, A. M., Ecker, M., Voit, W. E., Pancrazio, J. J. & Capadona, J. R. 2018. A Mosquito Inspired Strategy to Implant Microprobes into the Brain. Sci Rep, 8, 122.

Simon, D. M., Charkhkar, H., St John, C., Rajendran, S., Kang, T., Reit, R., Arreaga-SALAS, D., Mchail, D. G., Knaack, G. L., Sloan, A., Grasse, D., Dumas, T. C., Rennaker, R. L., Pancrazio, J. J. & Voit, W. E. 2017. Design and demonstration of an intracortical probe technology with tunable modulus. J Biomed Mater Res A, 105, 159–168.

Sohal, H. S., Jackson, A., Jackson, R., Clowry, G. J., Vassilevski, K., O’Neill, A. & Baker, S. N. 2014. The sinusoidal probe: a new approach to improve electrode longevity. Front Neuroeng, 7, 10.

Sridharan, A., Rajan, S. D. & Muthuswamy, J. 2013. Long-term changes in the material properties of brain tissue at the implant-tissue interface. J Neural Eng, 10, 066001.

Stevenson, I. H. & Kording, K. P. 2011. How advances in neural recording affect data analysis. Nat Neurosci, 14, 139–42.

Szarowski, D. H., Andersen, M. D., Retterer, S., Spence, A. J., Isaacson, M., Craighead, H. G., Turner, J. N. & Shain, W. 2003. Brain responses to micro-machined silicon devices. Brain Res, 983, 23–35.

Takeuchi, S., Ziegler, D., Yoshida, Y., Mabuchi, K. & Suzuki, T. 2005. Parylene flexible neural probes integrated with microfluidic channels. Lab Chip, 5, 519–23.

Toth, E., Fabo, D., Entz, L., Ulbert, I. & Eross, L. 2016. Intracranial neuronal ensemble recordings and analysis in epilepsy. J Neurosci Methods, 260, 261–9.

Vitale, F., Vercosa, D. G., Rodriguez, A. V., Pamulapati, S. S., Seibt, F., Lewis, E., Yan, J. S., Badhiwala, K., Adnan, M., Royer-Carfagni, G., Beierlein, M., Kemere, C., Pasquali, M. & Robinson, J. T. 2018. Fluidic Microactuation of Flexible Electrodes for Neural Recording. Nano Lett, 18, 326–335.

Weller, R. O., Sharp, M. M., Christodoulides, M., Carare, R. O. & Mollgard, K. 2018. The meninges as barriers and facilitators for the movement of Fluid, cells and pathogens related to the rodent and human CNS. Acta Neuropathol, 135, 363–385.

Wu, F., Tien, L. W., Chen, F. J., Berke, J. D., Kaplan, D. L. & Yoon, E. 2015. Silk-Backed Structural Optimization of High-Density Flexible Intracortical Neural Probes. Journal of Microelectromechanical Systems, 24, 62–69.

Xiang, Z., Yen, S.-C., Xue, N., Sun, T., Tsang, W. M., Zhang, S., Liao, L.-D., Thakor, N. V. & Lee, C. 2014. Ultra-thin flexible polyimide neural probe embedded in a dissolvable maltose-coated microneedle. Journal of Micromechanics and Microengineering, 24, 065015.

Xie, C., Liu, J., Dai, X., Zhou, W. & Lieber, C. M. 2015. Three-dimensional macroporous nanoelectronic networks as minimally invasive brain probes. Nature Materials, 14, 1286–92.

Zhao, Z., Li, X., He, F., Wei, X., Lin, S. & Xie, C. 2019. Parallel, minimally-invasive implantation of ultra-flexible neural electrode arrays. J Neural Eng.

Zhou, T., Hong, G., Fu, T. M., Yang, X., Schuhmann, T. G., Viveros, R. D. & Lieber, C. M. 2017. Syringe-injectable mesh electronics integrate seamlessly with minimal chronic immune response in the brain. Proc Natl Acad Sci U S A, 114, 5894–5899.

